# Human glial progenitor cells effectively remyelinate the demyelinated adult brain

**DOI:** 10.1101/822494

**Authors:** Martha Windrem, Steven Schanz, Lisa Zou, Devin Chandler-Militello, Nicholas J. Kuypers, John N. Mariani, Steven A. Goldman

**Affiliations:** Center for Translational Neuromedicine and the Department of Neurology, University of Rochester Medical Center, Rochester, NY 14642; Center for Translational Neuromedicine, University of Copenhagen, Denmark; Neuroscience Center, Rigshospitalet, Copenhagen, Denmark

**Keywords:** glial progenitor, oligodendrocytic progenitor, neural stem cell, demyelinating disease, cuprizone, leukodystrophy, multiple sclerosis, cell transplant

## Abstract

Human glial progenitor cells (hGPCs) can completely myelinate the brains of congenitally hypomyelinated shiverer mice, rescuing the phenotype and extending or normalizing the lifespan of these mice. We asked if implanted hGPCs might be similarly able to broadly disperse and remyelinate the diffusely and/or multicentrically-demyelinated *adult* CNS. In particular, we asked if fetal hGPCs could effectively remyelinate both congenitally hypomyelinated adult axons, *and* axons acutely demyelinated in adulthood, using adult *shiverer* mice and cuprizone-demyelinated mice, respectively. We found that hGPCs broadly infiltrate the adult CNS after callosal injection, and robustly myelinate congenitally-unmyelinated axons in adult *shiverer*. Moreover, implanted hGPCs similarly remyelinated denuded axons after cuprizone demyelination, whether they were delivered prior to *or* after initial cuprizone demyelination. Extraction and FACS of hGPCs from cuprizone-demyelinated brains in which they had been resident, followed by RNA-seq of the isolated human hGPCs, revealed their activation of transcriptional programs indicating their initiation of oligodendrocyte differentiation and myelination. These data indicate the ability of transplanted hGPCs to disperse throughout the adult CNS, to myelinate dysmyelinated regions encountered during their parenchymal colonization, and to also be recruited as myelinating oligodendrocytes at later points in life, upon demyelination-associated demand.

---

Oligodendrocytes are the sole source of myelin in the adult CNS, and their loss or dysfunction is at the heart of a wide variety of diseases of both children and adults. In children, the hereditary leukodystrophies accompany cerebral palsy as major sources of demyelination-associated neurological morbidity. In adults, demyelination contributes not only to diseases as diverse as multiple sclerosis and white matter stroke, but also to a broad variety of neurodegenerative and neuropsychiatric disorders (Lee et al., 2012; Roy et al., 2004; Tkachev et al., 2003). As a result, the demyelinating diseases have evolved as especially attractive targets for cell-based therapeutic strategies (Archer et al., 1997; Archer et al., 1994; Brustle et al., 1999; Yandava et al., 1999; Zhang and Duncan, 2000) (Franklin and Goldman, 2015; Goldman, 2016; Goldman et al., 2012). Prior studies have established the ability of neonatally-transplanted human glial progenitor cells (hGPCs) – also referred to interchangeably as oligodendrocyte progenitor cells (OPCs) or NG2 cells (Nishiyama et al., 2009) - to myelinate the hypomyelinated *shiverer* brain, and to rescue both the neurological phenotype and lifespan of neonatally-treated animals (Wang et al., 2013; Windrem et al., 2004; Windrem et al., 2008). Furthermore, a variety of enriched preparations of human GPCs, derived from both human brain and pluripotent stem cells, have been shown capable of remyelinating focal lesions of the adult brain and spinal cord (Buchet et al., 2011; Piao et al., 2015; Windrem et al., 2002). Yet these past studies were done in models of focal demyelination; it has remained unclear whether human GPCs are able to migrate extensively in adult brain tissue, as would be required for the repair of diffusely demyelinated tissue. Indeed, to our knowledge, no previous study has systematically assessed the ability of human GPCs to either migrate within or remyelinate brain tissue demyelinated in adulthood.

To that end, in this study we asked if fetal brain-derived hGPCs might exhibit sufficient migration and expansion competence in the adult environment to serve as therapeutic vectors for acquired adult-onset demyelination. In this regard, recent studies have supported the readiness with which axons can remyelinate after either congenital or focal adult demyelination, if provided myelinogenic cells (Buchet et al., 2011; Duncan et al., 2009; Ehrlich et al., 2017; Piao et al., 2015; Windrem et al., 2008). In that regard, recent reports have highlighted the cell-intrinsic loss of myelination competence as the basis for remyelination failure in disorders as diverse as progressive multiple sclerosis (Nicaise et al., 2017; Nicaise et al., 2019) and Huntington disease (Ferrari Bardile et al., 2019; Osipovitch et al., 2019), suggesting that the introduction of new, myelination-competent glial progenitors might be sufficient to achieve remyelination in those cases. On that basis, we asked here if human GPCs delivered directly into the adult brain could remyelinate axons in the setting of diffuse demyelination, as might be encountered clinically in multiple sclerosis and other causes of multicentric adult demyelination.

To do so, we used three distinct experimental paradigms: We first asked if hGPCs could effectively disperse within and myelinate the brains of *adult* MBP^*shi/shi*^ mice, which carry the *shiverer* mutation in the gene encoding myelin basic protein (MBP); this mutation precludes MBP expression and abrogates developmental myelination in these mice. By this means, we assessed the ability of hGPCs to restore myelin to the congenitally hypomyelinated adult brain, as might be encountered in the late postnatal treatment of a hypomyelinating leukodystrophy. Second, we next asked if *neonatally-engrafted* hGPCs could respond to cuprizone-induced *adult* demyelination by generating new oligodendrocytes and myelinating demyelinated axons, so as to assess the ability of already-resident hGPCs to remyelinate *previously-myelinated* axons, as might be demanded of human parenchymal progenitor cells after acquired demyelination. Third, we then asked if hGPCs transplanted into the *adult* brain, *after* cuprizone demyelination, could remyelinate denuded axons, as might be anticipated in the cell-based treatment of disorders such as progressive multiple sclerosis.

We found that in each of these experimental paradigms the hGPCs, whether engrafted neonatally or transplanted into adults, effectively dispersed throughout the forebrains, differentiated as oligodendroglia and myelinated demyelinated axons. These data suggested that transplanted hGPCs are competent to disperse broadly and differentiate as myelinogenic cells in the adult brain, and most critically, that they are able to remyelinate previously myelinated axons that have experienced myelin loss. On that basis, we also asked what the transcriptional concomitants of demyelination-associated mobilization might be in resident hGPCs. We therefore isolated hGPCs from neonatally-chimerized brains after the cessation of cuprizone demyelination, and used RNA-seq analysis to define those genes and cognate pathways induced by antecedent cuprizone demyelination. Together, these studies establish the ability of hGPCs to remyelinate demyelinated lesions of the adult human brain, while providing a promising set of molecular targets for the modulation of this process in human cells.

## RESULTS

### Adult shiverer mice exhibit myelination following hGPC delivery

To assess the ability of donor hGPCs to disperse and differentiate as oligodendroglia in the adult brain, we introduced CD140a-sorted fetal hGPCs into young adult shiverer x rag2^-/-^ immune-deficient mice (Sim et al., 2011b), as well as into two normally myelinated immunodeficient control lines, rag1^-/-^ on a C57Bl/6 background, and rag2^-/-^ on C3H. All mice were injected after weaning, over the range of 4-12 weeks of age; the shiverers were all injected between 4-6 weeks. A total of 22 mice (8 shiverers, 14 normally myelinated rag-null mice, both rag1^-./-^ and rag2^-./-^) were injected bilaterally in both the anterior and posterior corpus callosum, with 2 injections per hemisphere of 5 x 10^4^ hGPCs each. All 8 shiverers and 6 of the controls were sacrificed 12 - 15 weeks later at 19-22 weeks of age, while the remaining 8 control mice were sacrificed at approximately 1 year of age. The brains of all mice were examined for donor cell dispersal and oligodendrocytic differentiation as well as for MBP-immunoreactivity, which was necessarily donor-derived in the shiverer context.

The hGPCs proved both highly migratory and robustly myelinogenic in the adult brain. By 12-15 weeks after transplant, the injected cells had dispersed broadly throughout the forebrain, as is typically observed in similarly-treated neonates (Windrem et al., 2008), with a near-uniform distribution of donor cells noted throughout the white matter in both congenitally dysmyelinated shiverers (**Figure 1A**) and normally myelinated (**Figure 1B**) mice. Myelinogenesis was robust in the shiverers, with dense myelination of the corpus callosum (**Figures 1C-D**). Importantly, at the 19-week time-point assessed, the callosal densities of all human cells, as well as human GPCs, oligodendrocytes, and astrocytes, were all significantly and substantially higher in the recipient shiverer brains than in their myelin wild-type controls (**Figures 1E-H; 1I-K**), indicating the overwhelmingly competitive advantage of the human donor cells in the shiverer environment. In myelin wild-type control brains, whether examined at either 5 or 12 months, these cells also expanded and engrafted, but largely remained as progenitors (**Figure 1F**). These data indicated that CD140a-sorted hGPCs are able to migrate broadly throughout the young adult mouse brain, that the dispersal of these cells is not impeded by adult brain parenchyma, and that robust myelination of still-viable axons can begin even after a several months’ absence of mature myelin in the affected brain.

**Figure 1.**
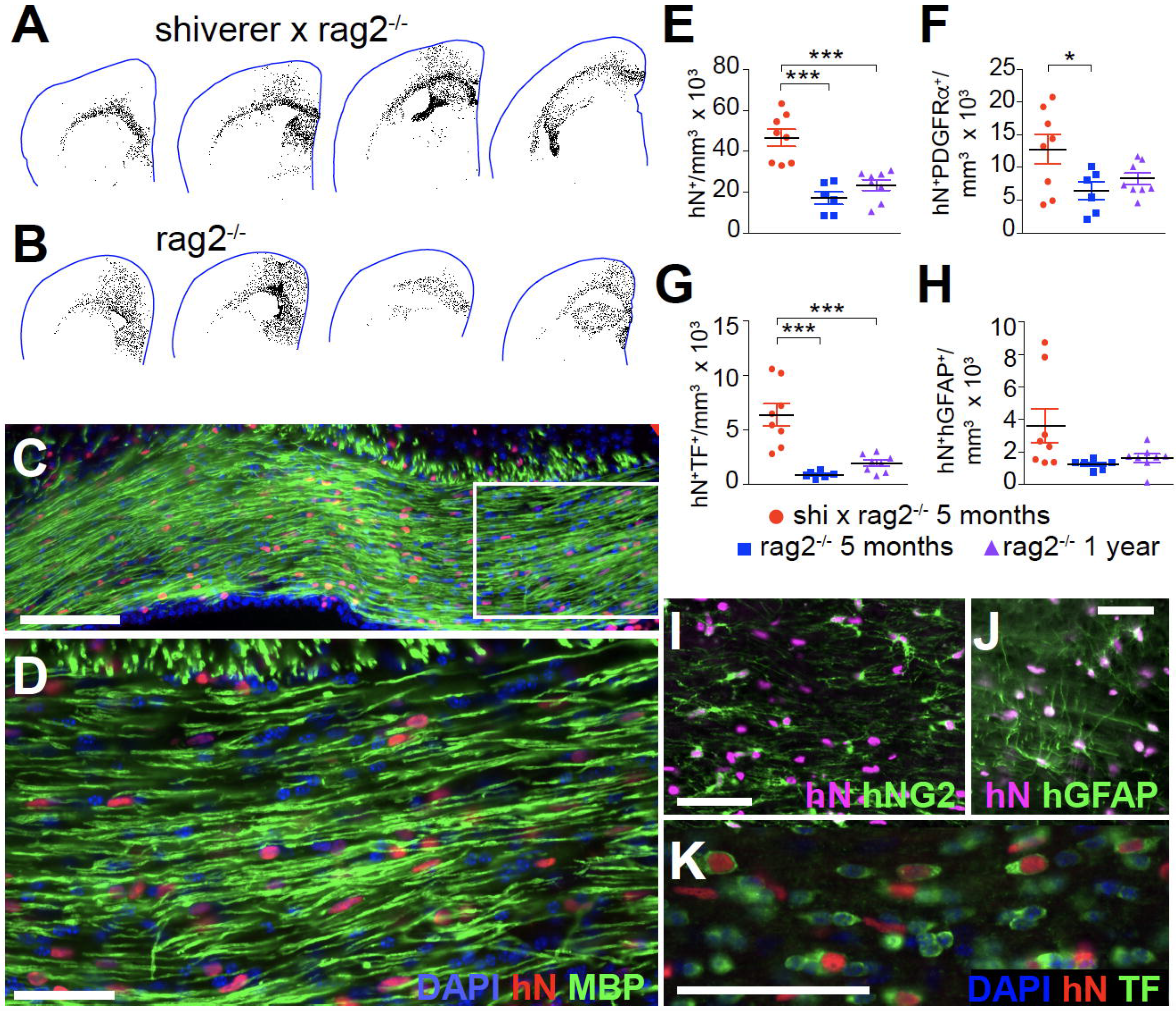
Human GPCs mediate robust myelination after transplantation into the adult shiverer brain. hGPCs proved both highly migratory and robustly myelinogenic, after delivery at 4-6 weeks of age to the hypomyelinated adult shiverer brain. **A**, By 19-20 weeks of age – 13-15 weeks after transplant - the injected cells had dispersed as broadly as is typically observed in similarly-transplanted shiverer neonates (Windrem et al., 2008, 2014), with a near-uniform distribution of donor cells noted throughout the forebrain white matter. **B**, hGPCs delivered to myelin wild-type rag2-/- mice distributed throughout both gray and white matter, though with less mitotic expansion than that noted in hypomyelinated shiverer recipients. **C**, Oligodendrocyte differentiation and myelinogenesis by donor hGPCs was robust, with dense myelination of the corpus callosum and fimbria. **D**, a higher power image of **C** shows the high proportion of donor cells in the now humanized host white matter. **E**, While both shiverer and myelin wild-type recipients exhibited substantial donor hGPC colonization after adult transplantation, the callosal densities of all human cells (**E**) and PDGFαR-defined hGPCs (**F**) were significantly higher in shiverer rather than myelin wild-type recipients. In the myelin wild-type recipients, the densities of all donor cells and identified hGPCs (**E-F**) were not significantly different between mice killed at 5 and 12 months of age, suggesting that donor cell expansion in the myelin wild-type brain occurred within the first 3 months after adult transplant, by 5 months of age. **G**, The density of transferrin (TF)-defined human oligodendroglia was 5-10-fold higher in adult-transplanted shiverers than in myelin wild-type hosts, when both were assessed 3 months after graft, at 5 months of age. **H**, A smaller proportion of the donor cell population matured as GFAP-defined astrocytes; these too proved significantly more abundant in the shiverer than wild-type hosts. **I-K**, representative images of anti-human NG2-defined donor-derived hGPCs (**I**), anti-human GFAP-defined astrocytes (**J**), and transferrin/human nuclear antigen co-expressing donor-derived oligodendrocytes (**K**) in 19-week old shiverer white matter, 13 weeks after transplantation at 6 weeks of age. Scale: **C-D:** 100 μm, **I-K**, 50 μm.

### Resident human GPCs can remyelinate the cuprizone-demyelinated corpus callosum

Cuprizone is a well-studied copper chelator, the chronic oral administration of which causes mitochondrial dysfunction that is both earliest and most prominent in myelinating oligodendrocytes (Morell et al., 1998). Its oral administration results in diffuse, relatively synchronous demyelination, which has been well-characterized in a variety of mouse strains and ages (Stidworthy et al., 2003). Cuprizone-induced demyelination is more reproducible than any other current model of demyelination, has little systemic toxicity at demyelinating doses, is associated with little acute axonal injury or neuronal loss, and is relatively non-inflammatory, except for local microglial activation (Matsushima and Morell, 2001). To assess the ability of human GPCs to remyelinate newly-demyelinated adult axons, we used dietary cuprizone to induce central demyelination, and followed the responses of both already-resident and later-introduced hGPCs to that myelin loss.

We began by asking if human GPCs *already resident* within the mouse white matter could differentiate as oligodendrocytes and remyelinate denuded axons after cuprizone challenge. We first assessed the effects of cuprizone on rag1^-/-^ x C57Bl/6 mice, and confirmed previous observations (Hibbits et al., 2009; Hibbits et al., 2012), that a 12-week course of cuprizone induced the widespread loss of transferrin (TF)-defined oligodendrocytes in the corpus callosum, with no detectable loss of resident mouse GPCs We then asked if neonatally-implanted *human* GPCs could similarly tolerate cuprizone exposure, and if so, whether they remained able to generate new oligodendrocytes and remyelinate adult-demyelinated axons. To that end, we transplanted rag1^-/-^ x C57Bl/6 mice on postnatal day 1 with human fetal A2B5^+^/PSA-NCAM^-^ GPCs, delivered as 10^5^ cells per hemisphere into the corpus callosum bilaterally. This protocol results in widespread colonization of the recipient brains by human GPCs, which ultimately replace many – and typically most - of the host murine GPCs (Windrem et al., 2014). The resultant human glial chimeric mice were then given dietary cuprizone (0.2% w/w) as a food additive, beginning at 4 months of age; by this time, the human NG2^+^ GPCs have largely replaced mouse callosal NG2^+^ cells (Windrem et al., 2014). The experimental mice were left on the cuprizone diet for 12 weeks, while littermate controls were maintained on a normal diet (**Figure 2A**). The density of human cells in the host white matter, as well as the percentage of those cells that differentiated as oligodendrocytes, was calculated for both cuprizone-treated and control mice at the start of the dietary manipulation, as well as at each of 3 different time-points: immediately after cuprizone cessation; after 8 weeks’ recovery; and after 20 weeks’ recovery - the latter out to 1 year of age.

**Figure 2.**
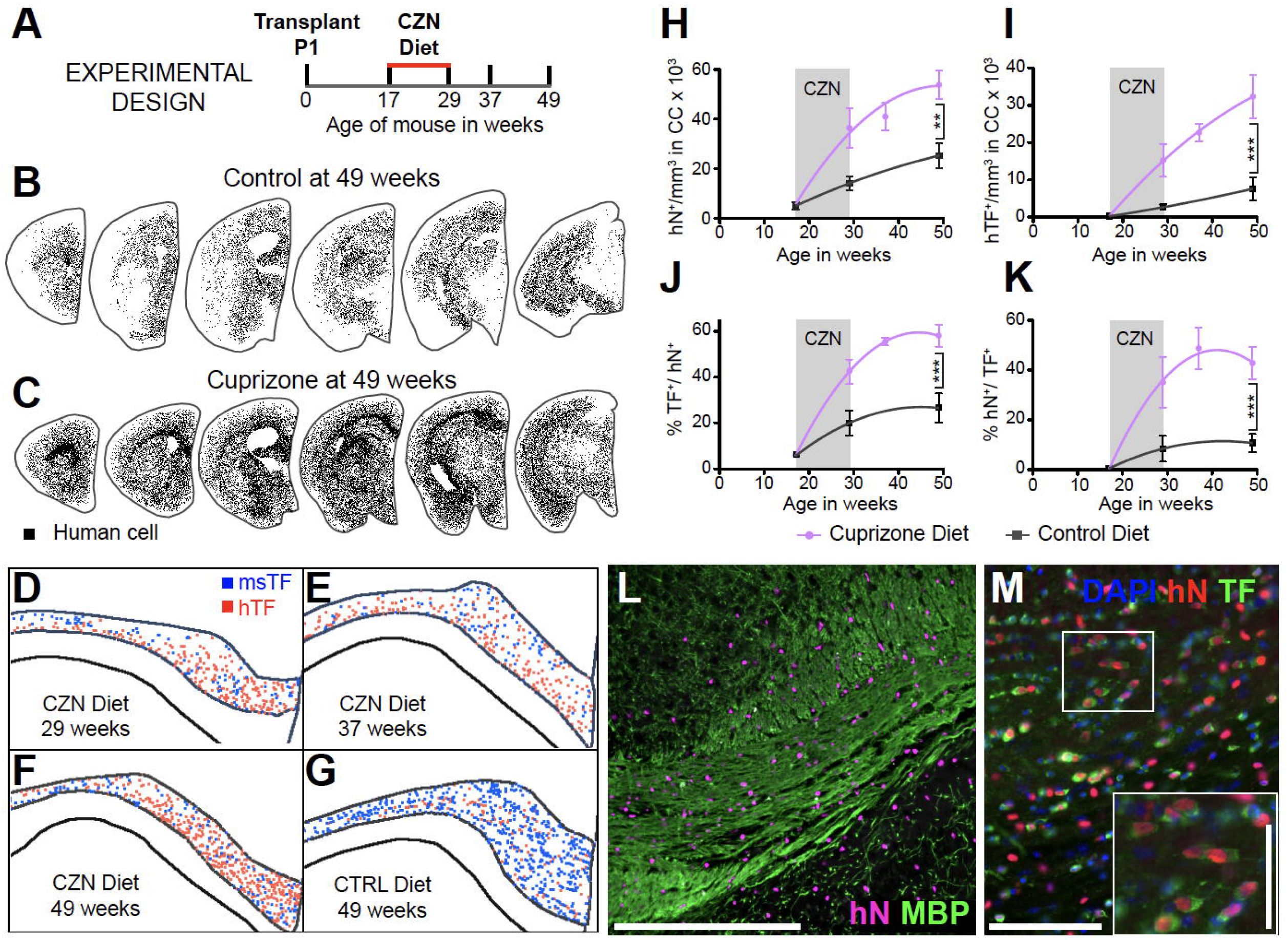
hGPCs differentiate as myelinogenic oligodendroglia in response to cuprizone demyelination. **A**, The schematic outlines the experimental design for neonatal engraftment followed by adult demyelination. Mice were transplanted with 2 x 10^5^ hGPCs perinatally, maintained on a control diet through 17 weeks of age, then placed on either a cuprizone-supplemented or normal diet for 12 weeks, then either sacrificed or returned to standard diet and killed at later time-points. **B-C.** Serial coronal sections comparing dot-mapped distributions of human (human nuclear antigen, hN) cells in control (**B**) and cuprizone-fed (**C**) mice at 49 weeks of age, following 20 weeks recovery on control diet. **D-G.** Relative positions and abundance of human (*red dots)* and mouse *(blue)* transferrin (TF)-defined oligodendrocytes, mapped in 20 μm coronal sections of corpus callosa of mice engrafted with hGPCs neonatally, demyelinated as adults from 17-29 weeks of age, then assessed either: **D**, at the end of the cuprizone diet; **E**, 8 weeks after return to control diet; or **F**, 20 weeks after cuprizone cessation. **G** shows an untreated control, age-matched to **F**. **H**, The density of human cells in the corpus callosum increases to a greater degree and more rapidly in cuprizone-demyelinated brains than in untreated controls, including during the 12 week period of cuprizone treatment *(indicated in gray)*. **I**, By 8 weeks after the termination of cuprizone exposure, the density of human oligodendroglia was >5-fold greater in cuprizone-demyelinated than untreated control brains. **J**, By that 8 week recovery point, over half of all hGPCs engrafted in the corpus callosa of cuprizone-treated mice had differentiated as oligodendrocytes, and accordingly (**K**), over half of all transferrin-defined callosal oligodendrocytes were human; in contrast, relatively few human oligodendrocytes were noted in untreated chimeric brains. **L**, Substantial colonization by human glia evident in this remyelinated corpus callosum, after 20 week recovery (human nuclear antigen, *magenta*; myelin basic protein, *green*). **M**, chimeric white matter populated, after cuprizone demyelination, by human GPC-derived oligodendroglia. Anti-human nuclear antigen (hNA) (*red*), transferrin, (*green*); inset highlights relative abundance of hNA^+^/transferrin^+^ human oligodendroglia. Scale: **L**, 100 μm; **M**, 50 μm, *inset*, 25 μm.

We found that the human GPCs tolerated cuprizone exposure at least as well as their mouse counterparts, and dispersed broadly throughout the forebrain (**Figures 2B-C**). The hGPCs then robustly generated new oligodendrocytes and effectively remyelinated the demyelinated white matter after cuprizone cessation (**Figures 2D-G; 2I-M**). In particular, both the total number of human cells, and the percentage that differentiated as oligodendrocytes, increased significantly faster and to a greater extent in the cuprizone-fed mice than in their matched controls (**Figures 2D-G**; also **2H-K**). The density of human cells in the corpus callosum of cuprizone-treated mice increased from 5,072 ± 1,611 at 4 months to 53,835 ± 5,898 cells/mm^3^ at one year; in contrast, over the same period, control mice exhibited a more modest expansion of GPCs, to 25,296 ± 4,959 cells/mm3; p=0.001 by 2-way ANOVA; F=7.40 (**Figure 2H**). Importantly, the proportion of all human cells that differentiated as mature oligodendrocytes by 1 year was twice that in the cuprizone-treated mice than in their untreated controls (58.0 ± 4.8% vs. 26.6 ± 6.4%; p<0.0001, F=13.32; **Figure 2J**). Similarly, the density of human oligodendrocytes rose over 4.5-fold in the cuprizone-treated mice, from 7,642 ± 3,095 to 32,323 ± 5,850 hTF^+^ cells/mm^3^ (p=0.0006; F=8.65; **Figure 2I**). These data indicate that cuprizone-induced demyelination yielded a relative increase in both the absolute numbers and relative proportions of parenchymal hGPC-derived oligodendrocytes, and the remyelination of demyelinated host axons by those cells (**Figures 2L-M**). Thus, those hGPCs already-resident within the callosal white matter responded to acute demyelination by differentiating as mature oligodendrocytes and remyelinating accessible denuded axons. Thus, resident hGPCs could myelinate not only axons that had never been myelinated, as in the adult shiverer brain, but also those that were previously ensheathed by myelin.

### Human GPCs can remyelinate axons when delivered after initial demyelination

We next asked if hGPCs delivered to the *adult* brain after initial demyelination and during ongoing cuprizone exposure, could migrate and myelinate host axons, and whether they were able to myelinate denuded axons as effectively as hGPCs resident in their host brains since neonatal development. To this end, we next used prolonged cuprizone exposure to demyelinate otherwise wildtype adult mice, and transplanted hGPCs into these demyelinating brains. In particular, to minimize the potential for endogenous remyelination by remaining mouse GPCs, we used a 20 week cuprizone course, which we found allowed much less spontaneous remyelination than shorter periods of cuprizone exposure (**Figure 3A**). We found that even when delivered into adult brain parenchyma 4 weeks after the onset of cuprizone treatment, during active demyelination, that the transplanted hGPCs not only dispersed widely, but did so and expanded more robustly than in untreated control brains (**Figures 3B-D; 3E-F**). When the cuprizone fed mice were assessed at 16 weeks after hGPC transplant (26 weeks of age), human oligodendrocytes were apparent, having differentiated from the engrafted hGPCs. By that point, more donor hGPCs had differentiated as oligodendrocytes in the cuprizone-demyelinated brains than in their untreated controls, suggesting both the preferential expansion of hGPCs (**Figure 3E**) and the active induction of oligodendrocytic phenotype by the demyelinated environment (**Figure 3F**). By 36 weeks post-transplant (46 weeks of age), allowing 20 additional weeks for phenotypic differentiation, over a quarter of all oligodendrocytes in the host white matter were of human origin (**Figure 3G**). Remarkably, the overall number and density of transferrin-defined oligodendrocytes, whether of mouse or human origin, was relatively preserved at all time points (**Figure 3H**). The transplanted human cells proceeded to robustly differentiate as oligodendrocytes and myelinate the demyelinated tissue, such that by 46 weeks of age - 36 weeks post-transplant - much if not most of the forebrain white matter in these previously cuprizone-demyelinated brains was of human origin (**Figures 3I-K**). Thus, human GPCs were able to effectively remyelinate mature axons that had been previously myelinated in the brain, and could do so even when delivered to the adult brain after the onset of demyelination.

**Figure 3.**
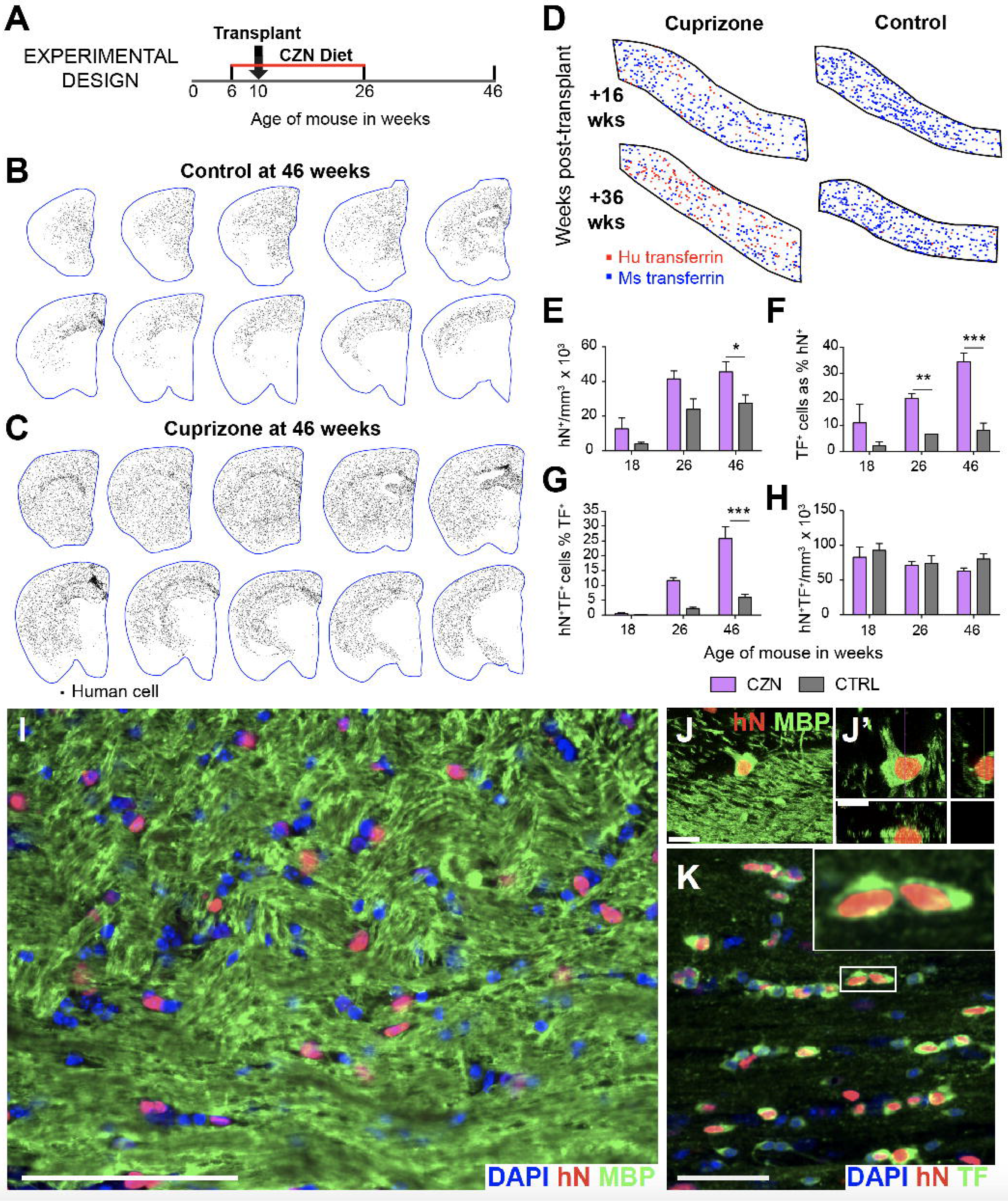
hGPCs differentiate and remyelinate axons after transplant into adult-demyelinated brain. **A**, At 6 weeks of age, experimental mice were put on a diet containing 0.2% cuprizone, while litter-mate controls remain on standard diet. At 10 weeks, 4 weeks into a 20 week cuprizone course, the mice were transplanted with 2 x 10^5^ hGPCs. Mice were sacrificed for histology either at the end of the cuprizone course (at 26 weeks) or after an additional 20-week recovery period (at 46 weeks). **B-C**, Maps show locations of individual human cells in 20 μm coronal hemi-sections of engrafted brains. **B**, Transplantation of hGPCs into a normally-myelinated 10-week old mouse yielded widespread engraftment, when mapped 36 weeks later at 46 weeks of age. **C**, In cuprizone-treated mice, transplanted hGPCs expanded to a significantly greater degree. **D-H**, Significantly more hGPCs differentiated as transferrin (TF)-defined oligodendrocytes in the cuprizone-demyelinated brains than in their untreated controls. **D**, hGPCs were more likely to differentiate as transferrin-expressing oligodendrocytes when transplanted into a demyelinating environment *(left)*, compared to a control brain *(right)*. **E-F**, The absolute density (**E**) and relative proportion (**F**) of human cells that differentiated as transferrin^+^ oligodendrocytes in the corpus callosum were respectively >5- and >10-fold greater in mice on the cuprizone diet than in their untreated controls. **G**, By 36 weeks posttransplant, over a quarter of all oligodendrocytes in the host white matter were of human origin. **H**, The overall density of transferrin-defined oligodendrocytes, whether of mouse or human origin, was relatively preserved at all time points. **I-K**, By 46-wks, adult-transplanted hGPCs are admixed with murine cells in the largely remyelinated corpus callosum (**I**), **J**, By this point, most myelinating oligodendrocytes in the cuprizone-demyelinated callosal were of human donor origin (human nuclear antigen, *red*; MBP, *green;* DAPI, *blue)*, just as many of the resident human cells had differentiated as TF-defined oligodendrocytes (**K**, human nuclear antigen, *red*; transferrin, *green*). Scale: **I**: 100 μm; **J**: 50 μm. K, 10 μm

### Human GPCs activated stereotypic transcriptional programs after cuprizone demyelination

We next asked whether demyelination and its attendant activation of human GPCs was associated with transcriptional events that might identify early determinants of progenitor cell mobilization, as well as those of astrocytic or oligodendrocytic fate. To thereby identify the responses of human GPCs to demyelination *in vivo*, we isolated them from cuprizone-demyelinated, neonatally-chimerized brains in which they had been resident, using CD140a-directed fluorescence-activated cell sorting, followed by RNA sequencing. To this end, neonatal *rag1^-/-^* mice were transplanted with fetal human hGPCs, and maintained through 12 weeks of age on a normal diet. At that point, control mice were continued on a normal diet, while experimental mice were transitioned to a diet of 0.2% (w/w) cuprizone for 12 weeks, to induce oligodendrocytic death. Cuprizone-demyelinated mice were then allowed to recover for an additional 12 weeks on a normal diet, before both groups were sacrificed at 36 weeks of age. The callosal white matter was then dissected, dissociated, and CD140a^+^ hGPCs isolated via FACS. The RNA of these hGPC isolates was then extracted and sequenced.

Principle component analysis of these normalized RNA-Seq samples revealed tight clustering of CTR samples, which as a group were readily distinguished from their post-CZN counterparts (**Figure 4A**). Both the post-CZN and CTR hGPCs were enriched for genes associated with early oligodendroglial lineage, including *CNP, GPR17, NKX2-2, OLIG1, OLIG2, SOX10, CSPG4, ST8SIA1*, as well as the selection marker *PDGFRA*/CD140a, the latter validating the selectivity and efficacy of the sort (**Figure 4B**). In contrast, the hGPC isolates exhibited low to undetectable expression of a number of neural stem cell, neural progenitor, endothelial, microglial, neuronal, and astrocytic markers. Yet while both groups presented with transcriptional signatures consistent with hGPC phenotype, a total of 914 transcripts were found to be differentially expressed between hGPCs following recovery from cuprizone treatment and their control counterparts (adjusted p<0.05). Of these, 777 genes were upregulated in cuprizone-treated GPCs, while 137 genes were down-regulated **(Supplementary Information, Table S1).** Functional analysis of this gene set demonstrated that the cuprizone-treated hGPCs differentially expressed gene ontologies reflecting cell proliferation, pathfinding and cell movement, and the initiation of myelination itself (**Figure 5**; and **Supplementary Information, Table S2**).

**Figure 4.**
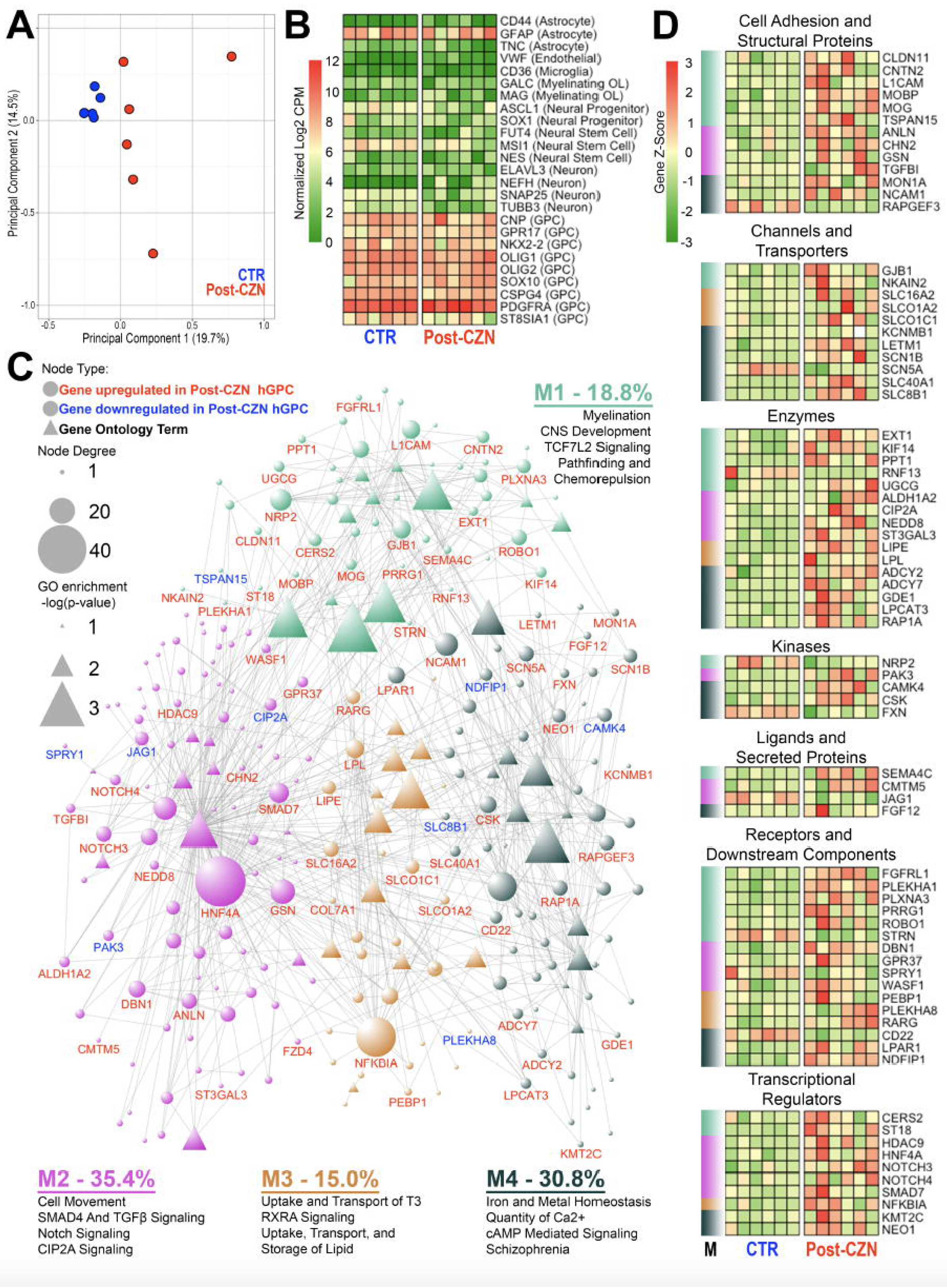
hGPC transcriptional networks augur compensatory remyelination after demyelination. Human glial chimeras were maintained on either a cuprizone (CZN)-containing or control diet from 12-24 weeks of age. 12 weeks later, at 36 weeks, the mice were killed and their resident hGPCs isolated via CD140a-based FACS, which were then subjected to RNA-Seq (n=6). **A**, Principle component analysis revealed tight clustering of hGPCs separated from post-CZN samples. **B**, Isolated hGPCs were enriched for genes indicative of an oligodendrocytic fate; gene expression representative of other lineages was minimal. **C**, A network was constructed from differentially expressed genes (*circles*) between post-CZN and CTR hGPCs (adjusted p<0.05); significantly associated gene ontology (GO) annotations (*triangles*) identified those pertinent and functionally related genes (*gene nodes*) that were differentially active in CZN-mobilized hGPCs. Gene node size was determined by the degree of connectivity, while annotation node sizes scaled with their adjusted p-values. Unsupervised modularity detection identified four modules (M) of closely related genes and annotations, for which a summary of annotations is provided along with the percentage of total gene connectivity for each module. Complete differentially expressed gene list is provided in the **Supplementary Information, Table S1**, while complete network information is offered in **Supplementary Information, Table S2. D**, A heatmap representation of genes identified in the previous GO network, organized by functional category and module membership (M).

**Figure 5.**
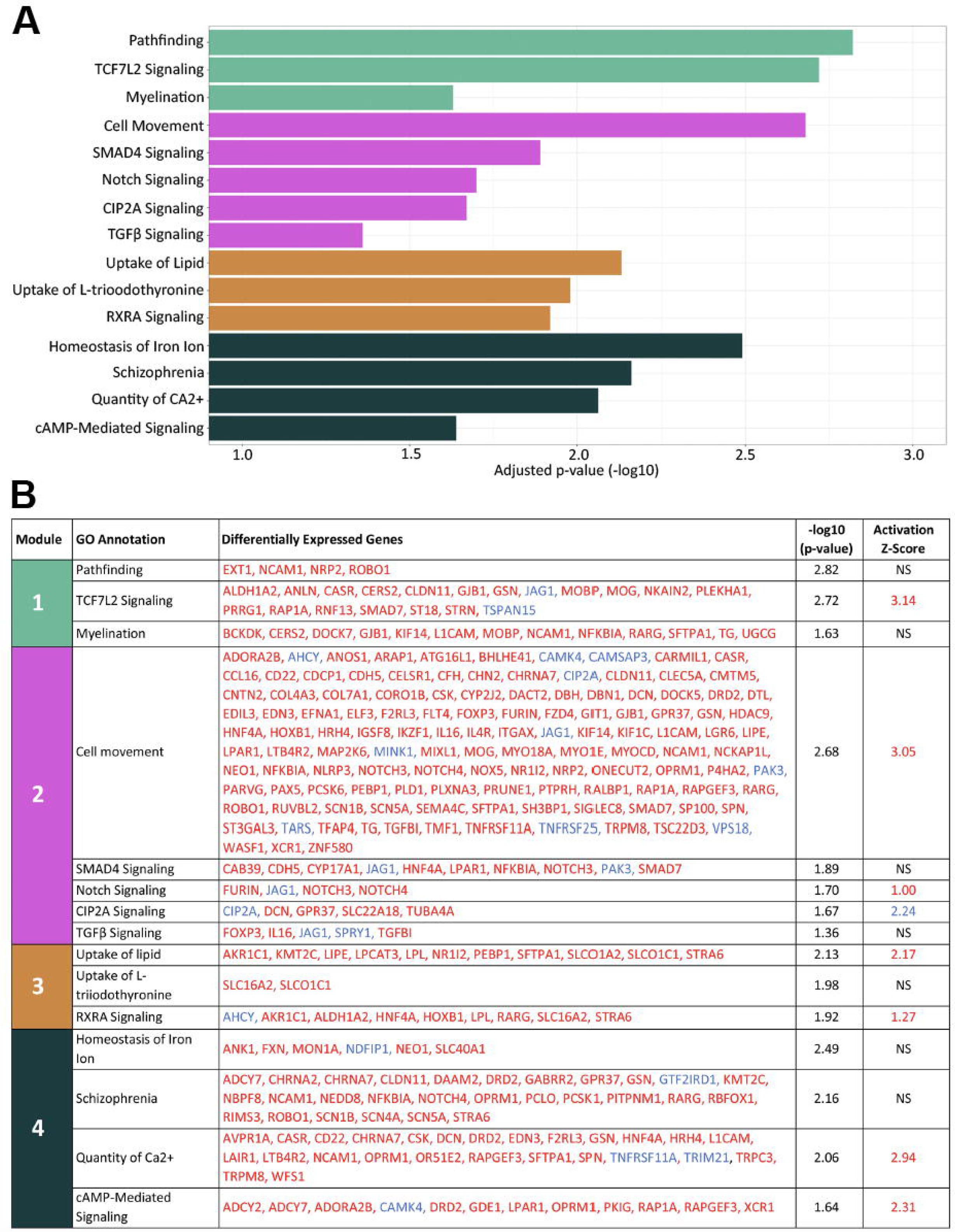
Enrichment of remyelination-associated pathways in cuprizone-exposed human GPCs. **A**, The significantly enriched functional categories highlighted in **Figure 4C** are organized here by color-defined modules. Enrichment was determined via Fisher’s Exact Test in Ingenuity Pathway Analysis. **B**, Genes differentially expressed by CZN-exposed hGPCs relative to controls, that contributed to these differentially-enriched pathways. Genes upregulated after cuprizone exposure in red, those down-regulated in blue. Activation Z-Scores are also provided for those pathways for which collective gene expression implies activation (*red*) or inhibition (*blue*), following CZN exposure in post-CZN vs. CTR hGPCs. Activation Z-Scores >1 were deemed significant.

### Network analysis revealed that cuprizone-exposed hGPCs were primed to oligoneogenesis

To aid in interpreting these data, a cuprizone-exposed hGPC expression network was constructed based upon both significantly enriched gene ontologies and differentially-expressed individual gene components thereof. The network included 43 significantly enriched and relevant functional terms, in addition to their contributing differentially expressed genes (network in **Figure 4C**; functionally-segregated heat-maps in **Figure 4D**; complete gene ontology network table in **Supplementary Information**, **Table S2**). Community detection via modularity analysis was then carried out to aggregate closely related functions and genes (Bastian et al., 2009; Blondel et al., 2008). This analysis yielded four distinct modules (M1-M4), which individually identified distinct processes associated with the initiation of remyelination by cuprizone demyelination-mobilized hGPCs.

M1 revealed that the hGPCs recovering from cuprizone demyelination markedly upregulated their expression of myelinogenesis-associated genes, including *MOG, MOBP*, and *CLDN11* (Goldman and Kuypers, 2015) (**Figure 4C**). Furthermore, several genes previously noted to be induced during oligodendrocyte differentiation and remyelination were also upregulated; these included *ST18* (Najm et al., 2013), *PLEKHA1* (Chen et al., 2015), and *CMTM5* (Doyle et al., 2008), along with those shown to be necessary for appropriate maturation of oligodendrocytes: *CERS2* (Imgrund et al., 2009), *LPAR1* (Garcia-Diaz et al., 2015), *GSN* (Zuchero et al., 2015), *KIF14* (Fujikura et al., 2013), *CNTN2* (Zoupi et al., 2018), *ST3GAL3 (Yoo et al., 2015)*, and *WASF1/WAVE1* (Kim et al., 2006). Interestingly, M1 further revealed that TCF7L2 signaling, a major driver of myelination (Hammond et al., 2015; Zhao et al., 2016), is also strongly upregulated in remyelinating hGPCs Genes contributing to the migration and pathfinding of GPCs were also up-regulated following cuprizone exposure; these included *ROBO1* (Liu et al., 2012) and the class 3 semaphorin co-receptors *NRP2* (Boyd et al., 2013) and *PLXNA3* (Xiang et al., 2012).

Within M2, a number of transcription factors and associated signal effectors vital to oligodendrocyte movement and differentiation were noted to be significantly enriched in the post-cuprizone hGPCs. These included SMAD4 (Choe et al., 2014), TGFβ (McKinnon et al., 1993), and NOTCH (Park and Appel, 2003), as well as the pro-myelinogenic genes *SMAD7* (Weng et al., 2012), *PAK3* (Maglorius Renkilaraj et al., 2017), and *NOTCH3* (Zaucker et al., 2013). In contrast, the Notch pathway inhibitor of differentiation *JAG1* was sharply repressed (John et al., 2002), suggesting the incipient differentiation of these cells. Also localizing to this module and upregulated in remyelinating GPCs was *GPR37*, the expression of which attends and is necessary for oligodendrocyte differentiation (Smith et al., 2017; Yang et al., 2016). Interestingly, *CIP2A*, a transcriptional repressor of *GPR37*, was profoundly down-regulated in cuprizone-mobilized hGPCs, again suggesting that cuprizone-demyelination triggers the active disinhibition of oligodendrocytic differentiation by previously quiescent parenchymal hGPCs (Yang et al., 2016).

M3 consisted of several functional categories strongly involved in myelination. These included transcripts involved in the transport and uptake of thyroid hormone and L-triiodothyronine (Almazan et al., 1985; Bhat et al., 1979). M3 also included genes associated with retinoid-signaling, particularly RXRA and the retinoid receptor complex partner *RARG*, the up-regulation of which was observed in hGPCs after cuprizone treatment (Huang et al., 2011; Tomaru et al., 2009). This is of particular significance as this signaling family has previously been tied not only to developmental myelination (de la Fuente et al., 2015), but also to remyelination as well (Huang et al., 2011). M3 also included genes associated with cholesterol and lipid uptake, processes critical to myelination (Saher et al., 2005).

The fourth module included genes associated with the transport and homeostatic regulation of iron and other multivalent cations, which were upregulated following cuprizone demyelination. Iron in particular has been reported to be important in the regulation of oligodendrocytic differentiation and myelination (Connor and Fine, 1987; Morath and Mayer-Prschel, 2001). In this regard, upregulation of *MON1A* and *FXN*, iron metabolism-associated genes strongly dysregulated in MS lesions (Hametner et al., 2013), was observed in cuprizone-mobilized hGPCs. Similarly, we noted that genes encoding calcium regulatory proteins associated with myelin maturation, which included *CASR, GSN*, and *TRPC3* (Cheli et al., 2015; Krasnow et al., 2018), were also increased in hGPCs after cuprizone demyelination, as were transcripts involved in cAMP signaling, another modulator of oligodendrocyte differentiation, in part via crosstalk with GPR37 and GPR17 (Simon et al., 2016; Yang et al., 2016).

Overall, the pattern of differential gene expression by cuprizone-exposed hGPCs reflected in **Figure 4** appears to define an expression network typifying that of early progenitor-derived remyelination. As such, these data indicate that when mobilized in response to antecedent cuprizone demyelination, human GPCs activated a coherent set of transcriptional programs that served to direct both oligodendrocytic differentiation and myelinogenesis.

## DISCUSSION

The congenitally hypomyelinated shiverer mouse (*MBP^shi/shi^*) is a naturally-occurring mutant that lacks myelin basic protein (MBP), and as such cannot make compact myelin. We have found that the intracerebral injection of hGPCs into neonatal shiverer mice results in the widespread dispersal of the human donor cells, followed by their oligodendrocytic differentiation and myelinogenesis (Windrem et al., 2004; Windrem et al., 2008). This ultimately leads to the complete or near-complete myelination of the recipient’s brain, brainstem and spinal cord, attended by the clinical rescue of a large proportion of transplanted neonates. Yet notwithstanding the robust developmental myelination of the host CNS by neonatally-delivered hGPCs, in order for hGPC delivery to be a viable regenerative strategy for treating adult demyelination - especially as occurs in multicentric and diffuse myelin loss - then the donor cells must be capable of migrating within and myelinating *adult* brain parenchyma.

In adults, oligodendrocytic loss contributes to diseases as diverse as hypertensive and diabetic white matter loss, traumatic spinal cord and brain injury, multiple sclerosis (MS) and its variants, and even the age-related white matter loss of the subcortical dementias. All of these conditions are potential targets of glial progenitor cell replacement therapy, recognizing that the adult disease environment may limit this approach in a disease-specific fashion (Goldman, 2017; Goldman et al., 2012). For instance, the chronically ischemic brain tissue of diabetics with small vessel disease may require aggressive treatment of the underlying vascular insufficiency before any cell replacement strategy may be considered. Similarly, the inflammatory disease environments of multiple sclerosis as well as many of the leukodystrophies present their own challenges, which need to be overcome before cell-based remyelination can succeed (Franklin and ffrench-Constant, 2008; Franklin and Ffrench-Constant, 2017; Goldman, 2016; Ip et al., 2006); these include participation by mobilized GPCs in the inflammatory response, potentially complicating therapeutic efforts further (Falcao et al., 2018). Nonetheless, current disease-modifying strategies for treating both vascular and autoimmune diseases have advanced to the point where stabilization of the disease environment can often be accomplished, such that transplant-based remyelination for the structural repair of demyelinated adult white matter may now be feasible.

Despite concerns as to the ability of glial progenitor cells to remyelinate axons in disease environments such as those associated with multiple sclerosis and the periventricular leukomalacia of cerebral palsy, a number of studies have pointed to the cell-intrinsic nature of oligodendrocytic differentiation block in these cases. These studies have suggested that the inability of parenchymal glial progenitors to produce myelinating oligodendrocytes in these conditions is a function of stable epigenetic blocks in the differentiation potential of these cells, imparted by the specific disease process or its antecedents. Newly introduced naïve hGPCs might thus be expected to exercise unfettered differentiation and myelination competence in host brains, and as such be able to remyelinate previously demyelinated axons. Indeed, several prior studies have indicated the ability of transplanted oligodendrocyte progenitors to remyelinate adult-demyelinated central axons (Duncan et al., 2009; Mozafari et al., 2015). To define the competence of human GPCs to remyelinate axons when delivered to the demyelinated adult brain, we used two different antigenic phenotypes of GPCs, respectively defined as CD140a^+^ and A2B5^+^/PSA-NCAM^-^, each derived from fetal human brain tissue (Sim et al., 2011a; Windrem et al., 2008). These antigenic phenotypes are largely but not completely homologous; the CD140a phenotype is the major fraction of, and largely subsumed within, the A2B5^+^/PSA-NCAM^-^ pool (Sim et al., 2011a). We assessed the dispersal and myelination competence of these cell types in two distinct adult models of myelin deficiency, the congenitally hypomyelinated shiverer mouse as well as the normally myelinated adult mouse, and the cuprizone-treated demyelinated adult mouse.

In each of these model systems, the transplanted hGPCs effectively restored myelin to the host brain. Donor-derived myelination was robust when cells were delivered to the adult shiverer brain, just as we had previously reported after hGPC transplants in neonatal shiverer mice. The design of this experiment was intended to mimic what might be encountered in the postnatal treatment of a hypomyelinating leukodystrophy, and the effective progenitor cell dispersal and myelination that we observed augurs well for the potential of this approach in the treatment of children with congenital leukodystrophies. Similarly, donor hGPC-derived oligodendrocyte differentiation and axonal remyelination proved robust in response to cuprizone demyelination, whether by hGPCs already resident within the adult-demyelinated brains, or by those transplanted during and after demyelination. Together, this latter set of experiments in particular provided an important proof-of-principle, showing that hGPCs could remyelinate axons that had already been myelinated in the past, and which were then demyelinated in the setting of oligodendrocyte loss; precisely such a scenario might be anticipated in disorders such as progressive multiple sclerosis, in which the remyelination of stably-denuded residual axons might be expected to confer functional benefit.

Indeed, in each of these experimental paradigms we found that the hGPCs, whether engrafted neonatally or transplanted into adults, effectively dispersed throughout the forebrains, even in normally myelinated mice, and differentiated as oligodendroglia and myelinated demyelinated lesions as these evolved or were encountered. These data suggest that transplanted fetal hGPCs are competent to disperse broadly and differentiate as myelinogenic cells in the adult brain, and – critically-that they are able to remyelinate previously myelinated axons that have experienced myelin loss.

Having established the potential of human GPCs as myelinogenic vectors, we then used RNA-seq of hGPCs extracted from the brains in which they had been resident during cuprizone exposure, to assess the transcriptional response of these cells to demyelination and initial remyelination. By this means, we correlated the demyelination-associated recruitment of resident hGPCs with their coincident transcriptional responses, so as to identify - in human cells, isolated directly from the in vivo environment - those genes and pathways whose targeting might permit the therapeutic modulation of both progenitor recruitment and differentiated fate.

We found that mobilized human GPCs indeed expressed a transcriptional signature consistent with early remyelination, as has been noted with demyelination-mobilized murine progenitors as well (Moyon et al., 2015). Yet human and mouse glial progenitors are quite different phenotypes, and are distinct in their transcriptional signatures (Lovatt et al., 2007; Sim et al., 2006; Sim et al., 2009; Zhang et al., 2016), lineage restriction (Nunes et al., 2003), and daughter cell morphologies (Oberheim et al., 2009). Accordingly, we noted a number of activated pathways in demyelination-triggered hGPCs not previously noted in models of murine demyelination. These included a number of upstream directors of oligodendroglial fate choice and differentiation, that included TCF7L2, TGFß/SMAD4, and NOTCH-driven pathways. The activation of each of these pathways was linked to the demyelination-associated disinhibition of differentiation by these parenchymal hGPCs, the oligodendrocytic maturation of which then enabled the compensatory remyelination of denuded axons. In addition, besides the activation of differentiation-associated pathways in demyelination-stimulated hGPCs, our data also suggested the critical importance during early remyelination of pathways enabling iron transport and metabolism, as well as those facilitating cholesterol and lipid uptake, pathways that are critically important to oligodendrocytic myelinogenesis. Interestingly, the identification of these pathways as differentially upregulated during remyelination suggests that the efficiencies of both myelinogenesis and myelin maturation might be further potentiated via metabolic optimization, and potentially even by dietary modulation referable to these pathways. Together, these studies establish an operational rationale for assessing the ability of hGPCs to remyelinate demyelinated lesions of the adult human brain, while providing a promising set of molecular targets for the modulation of this process in human cells.

## STAR METHODS

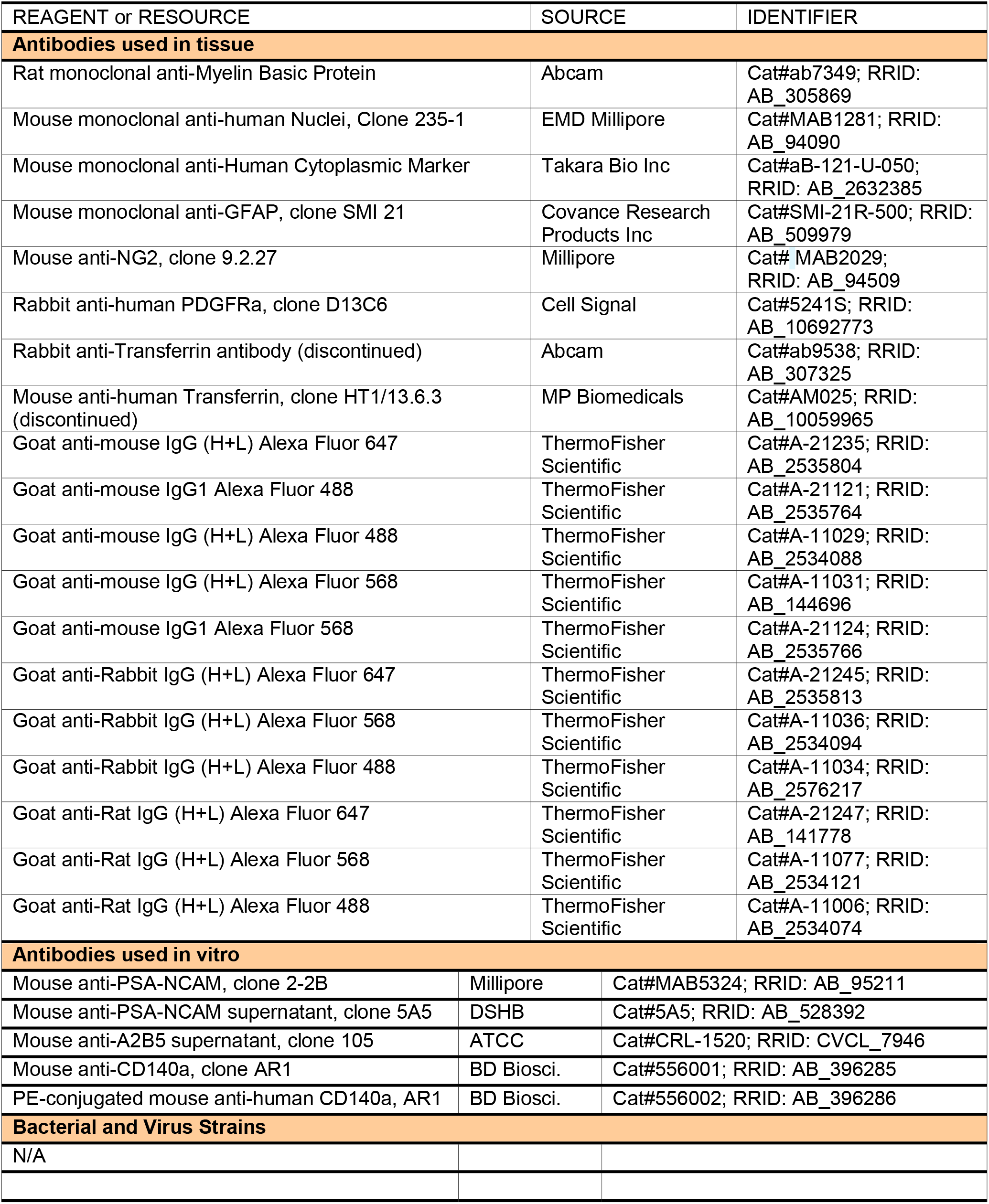
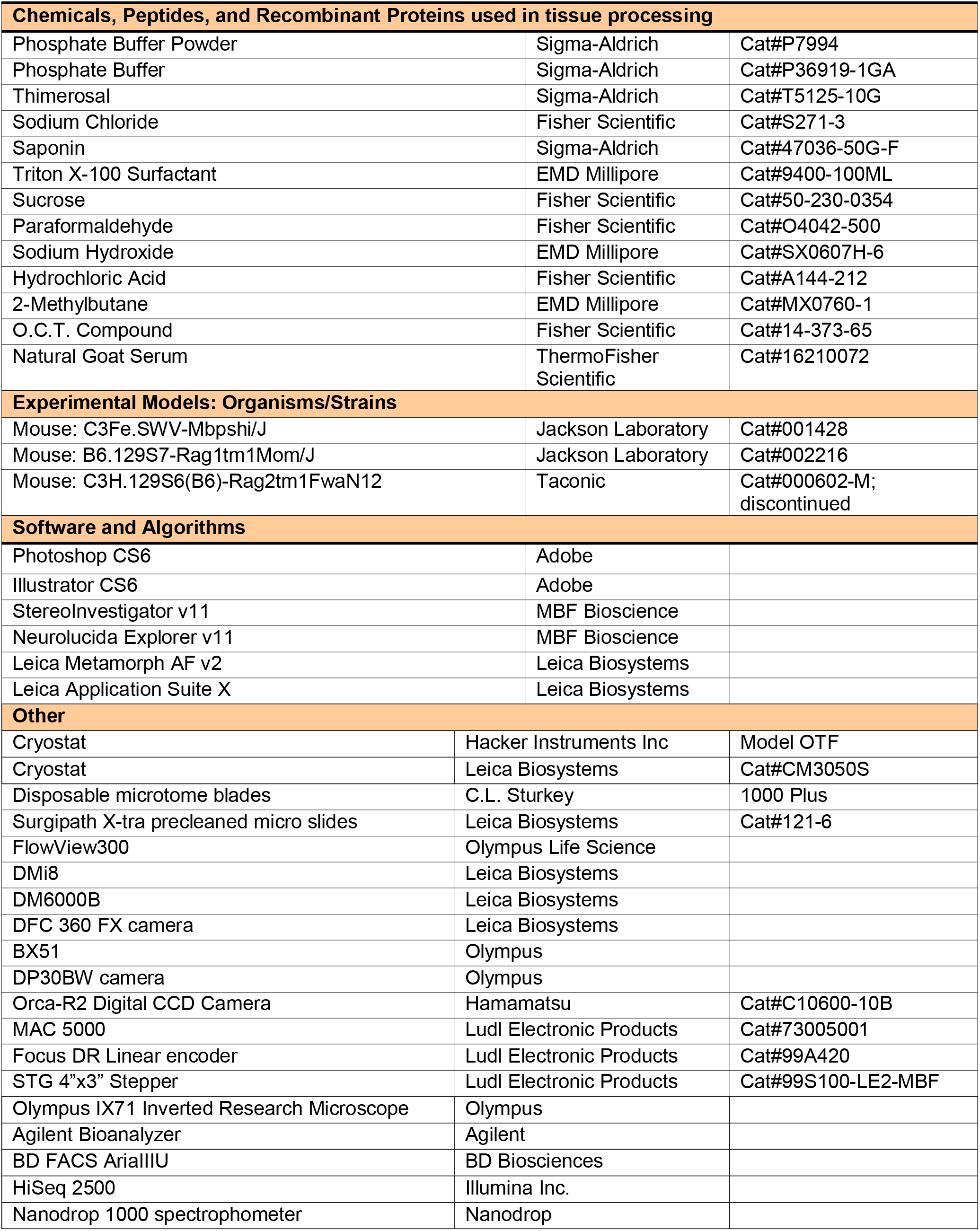
KEY RESOURCES TABLE.

### Lead Contact and Materials Availability

Further information and requests for resources and reagents should be directed to and will be fulfilled by the Lead Contact, Steven A. Goldman.

### Experimental Model and Subject Details

#### Animal models and transplantation

##### Shiverer mice, engrafted as adults

Homozygous Shi^-/-^ x rag2^-/-^, rag2^-/-^, and rag1^-/-^ immunodeficient mice were bred and housed in a pathogen-free environment in accordance with University of Rochester animal welfare regulations. Mice from each genotype were transplanted between the ages of 4-12 weeks with 1 x 10^5^ hGPCs/1 μl/hemisphere (n=2-4 mice/time-point/genotype), delivered bilaterally to the genu of the corpus callosum at coordinates: AP −0.8; ML ± 0.75; DV −1.25, all relative to bregma. Mice were injected with FK506 (5 mg/kg, i.p.; Tecoland, Inc.) daily for 3 days pre- and 3 days post-surgery. All shi^-/-^ x rag2^-/-^ mice were killed at age 19-20 weeks, or when clinical morbidity, as defined in our animal welfare policy, was observed. For the myelin wildtype mice, half of all rag2^-/-^ and rag1^-/-^ animals were sacrificed between 20-22 weeks of age, and the other half at 1 year.

##### Myelin wild-type mice, neonatally transplanted, cuprizone-demyelinated as adults

Homozygous rag1-null immunodeficient (rag1^-/-^) mice on a C57BL/6 background were bred in our colony. Animals were transplanted with hGPCs neonatally, via bilateral injections delivered to the presumptive corpus callosum (Windrem et al., 2004), so as to engraft newborn recipient brains before cuprizone demyelination. Beginning at 17 weeks of age, these mice were fed *ad libitum* a diet containing 0.2% (w/w) cuprizone (S5891, BioServe) for 12 weeks and then returned to normal diet. Littermate and non-littermate controls were maintained on a normal diet. Mice were sacrificed before diet (17 weeks), during diet (25 weeks), immediately after diet completion (29 weeks), and after either 8 weeks (37 weeks old) or 20 weeks (49 weeks old) of post-cuprizone recovery.

##### Myelin wild-type mice, cuprizone-demyelinated as adults, then transplanted

Homozygous rag1-null mice were subjected to cuprizone demyelination as noted, for a 20-week period beginning at 6 weeks of age. They were transplanted with hGPCs at 10 weeks of age, 4 weeks into their period of cuprizone demyelination. At that point, the mice were transplanted with a total of 200,000 PSA-NCAM^-^/A2B5^+^ cells, delivered sterotaxically as 1×10^5^ hGPCs/1 μl HBSS into the corpus callosum bilaterally at the following coordinates: from bregma, AP −0.8 mm, ML ±0.75; from dura, DV - 1.25 mm. Upon recovery, mice were returned to their cages. Mice were injected with FK506 (5 mg/kg, i.p.; Tecoland, Inc.) daily for 3 days pre- and 3 days post-surgery. Mice were sacrificed during diet (18 weeks), immediately after diet completion (26 weeks), and after 20 weeks (46 weeks old) of post-cuprizone recovery.

#### Cells

Human glial progenitor cells (hGPCs) were sorted from 18-22 week g.a. human fetuses, obtained from the surgical pathology suite, by either A2B5- or CD140a-directed isolation. Acquisition, dissociation and immunomagnetic sorting of A2B5^+^/PSA-NCAM^-^ cells were as described (Windrem et al., 2004). GPCs were isolated from dissociated tissue using a dual immunomagnetic sorting strategy: depleting mouse anti-PSA-NCAM^+^ (Millipore, DSHB) cells, using microbead tagged rat anti-mouse IgM (Miltenyi Biotech), then selecting A2B5^+^ (clone 105; ATCC, Manassas, VA) cells from the PSA-NCAM^-^ pool, as described (Windrem et al., 2004; Windrem et al., 2008). After sorting, cells were maintained for 1-14 days in DMEM-F12/N1 with 10 ng/ml bFGF and 20 ng/ml PDGF-AA. Alternatively for some experiments, CD140a/PDGFαR-defined GPCs were isolated and sorted using MACS as described (Sim et al., 2011b), yielding an enriched population of CD140^+^ glial progenitor cells.

### Methods Details

#### Histology

All mice were perfused with HBSS (-) followed by 4% paraformaldehyde. Brains were cryopreserved with 6%, then 30% sucrose and embedded coronally in OCT (TissueTek). Brains were then cut at 20 μm on a Leica cryostat. Sections were processed for one or more of the antigenic markers (see Table below).

#### Quantification

The optical fractionator method was used to quantify the phenotype of cells in the corpus callosum for: oligodendrocytes (transferrin), astrocytes (GFAP), and progenitors (PDGFRα). Transferrin, a cytoplasmic and membrane-localized iron transport protein, permits identification of colabeling with human nuclear antigen for the purpose of quantification by species of origin (Connor and Fine, 1987; Connor et al., 1993). Quantification of the phenotypes in the corpus callosum was performed by using a computerized stereology system consisting of a BX-51 microscope (Olympus) equipped with a Ludl (Hawthorne, NY) XYZ motorized stage, Heidenhain (Plymouth, MN) z-axis encoder, an Optronics (East Muskogee, OK) QuantiFire black and white video camera, a Dell (Round Rock, TX) PC workstation, and Stereo Investigator software (MicroBrightField, Wiliston, VT). Within each corpus callosum, beginning at a random starting point where it crosses the midline, 3 sections equidistantly spaced 480 μm apart were selected for analysis. The corpus callosum was outlined from the midline to 1 mm lateral, and all cells were counted. Upper and lower exclusion zones of 10% of section thickness were used.

#### RNA-Sequencing and analysis of FACS isolated hGPCs

To observe transcriptional changes in hGPCs following demyelination neonatal *rag1^/-^* mice were transplanted with 2 x 10^5^ hGPCs as described above. At 12 weeks, mice either maintained a normal diet or were transitioned to a 0.2% (w/w) cuprizone diet fed *ad libitum* until 24 weeks of age when they were returned to a normal diet. At 36 weeks, all mice were anesthetized with sodium pentobarbital, perfused with ice cold HBSS^+/+^ (containing calcium and magnesium) (Thermo Fisher), and the brains extracted and placed into fresh HBSS^+/+^ in a tissue culture plate. Excess HBSS^+/+^ was removed and the corpus callosum of each animal was isolated surgically, minced, and suspended in HBSS^-/-^. Corpora callosa from two mice from the same experimental group were pooled for each study. The tissue was transferred to a 15 ml conical tube and rinsed twice with HBSS^-/-^. The tube was filled to maximum volume with HBSS-/-, and then centrifuged for 5 min at 1,500 rpm. The supernatant was then removed and 0.62 units Research Grade Liberase DH (Roche) was added, and the samples then incubated at 37°C for 40 min with gentle rocking. Following Liberase dissociation, 2 ml 0.5% BSA in MEM (Thermo Fisher) supplemented with 500 units bovine pancreas DNase (Sigma) and 0.5 ml PD-FBS (Cocalico Biologicals) was added to the sample. The sample was triturated with a p1000 Pipetman and then passed through a 70 μm cell strainer. MEM containing 0.5% BSA was then added to the tube to bring it to full volume, and the tube centrifuged at 1,500 rpm for 10 min. Dissociated cells were then tagged with anti-CD140a-PE and sorted via FACS as previously reported (Sim et al., 2011b). Cells were lysed and prepared for library construction via Prelude Direct Lysis Module (NuGEN) according to the manufacturer’s protocol.

Libraries were constructed using Ovation RNA-Seq System V2 (NuGEN) according to manufacturer’s protocol and sequenced with a read length of paired-end 125bp on a HiSeq 2500 system (Illumina). Reads were demultiplexed and cleaned using Trimmomatic (Bolger et al., 2014). Reads were aligned to human genome GRCh38,p10 and mapped to Ensembl reference 91 via STAR 2.5.2b (Dobin et al., 2013), with quantMode set to TranscriptomeSAM. Gene abundances and expected counts were then calculated using RSEM 1.3.0 (Li and Dewey, 2011). Expected counts were imported into R via tximport for differential expression analysis between cuprizone and control hGPCs (R Core Team, 2017; Soneson et al., 2015). Low-expressing genes were removed prior to analysis if their expected counts fell below a median of 3 in both conditions. Full within-lane normalization of samples was conducted using EDASeq to adjust for GC-content effects prior to the generation of library size adjusted counts via DESeq2 (Love et al., 2014; Risso et al., 2011). Differential expression between post-cuprizone and control hGPCs was then determined using DESeq2 following the addition of a variance factor to the generalized linear model. This factor was calculated using RUVSeq’s RUVs function with all genes set as *in silico* negative controls (Risso et al., 2014). Genes with an adjusted p-value <0.05 were considered significant. Only genes with mean transcripts per million (TPM) >6.5 in either group were kept for functional analysis, as they were more likely to be biologically significant.

For functional analysis and inference of gene interactivity, a gene ontology network was constructed. Differentially expressed genes between both groups were analyzed in Ingenuity Pathway Analysis (QIAGEN) where 43 significantly enriched terms were selected based on relevance, along with their contributing differentially expressed genes, to be used as nodes. Along with the undirected edges derived from gene-GO term associations, edges were further generated via IPA’s curated database connecting genes with known interactions. Network visualization was carried out in Cytoscape (Shannon, 2003) with the determination of modularity occurring in Gephi (Bastian et al., 2009). Nodes were clustered within their respective modules and aesthetically repositioned slightly. Gene expression data are available via GEO, accession number GSE112557.

##### Antibodies

Phenotyping of donor cells was accomplished by immunostaining for human nuclear antigen (Millipore, clone MAB1281), together with one or more of the following:

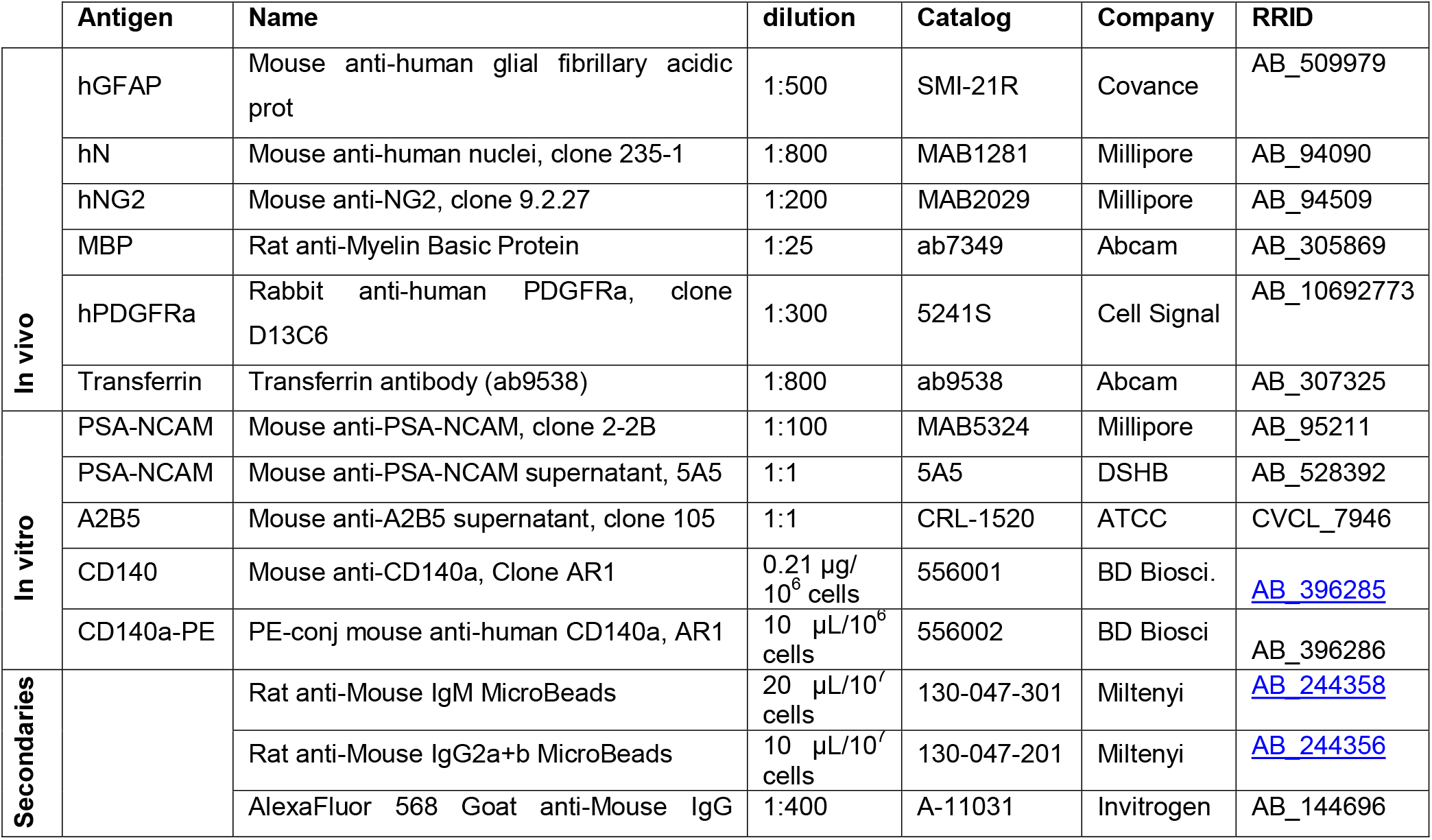
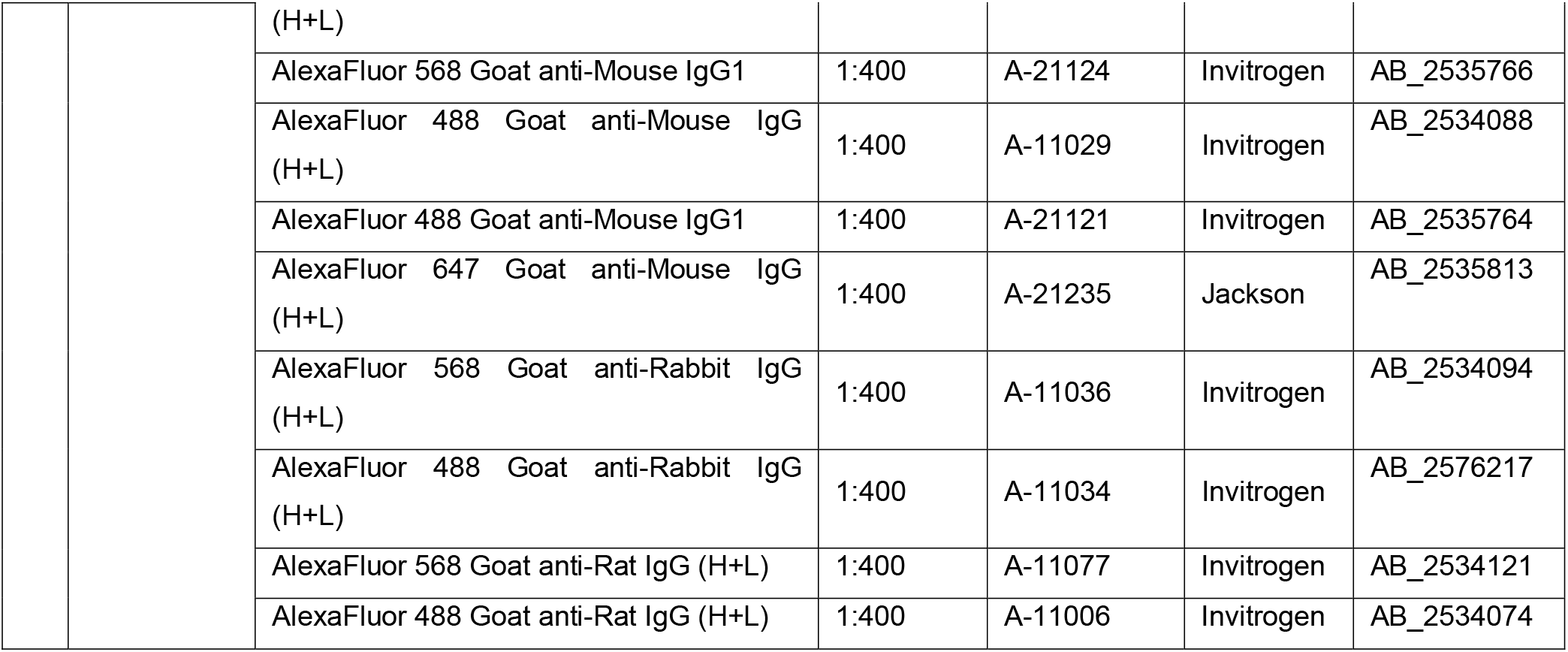

## Acknowledgements

Support by NINDS grants R01NS110776 and R01NS75345, the Dr. Miriam and Sheldon G. Adelson Medical Research Foundation, the Mathers Charitable Foundation, the NY Stem Cell Research Program (NYSTEM), the Oscine Corporation, Sana Biotechnology, the Novo Nordisk Foundation, and the Lundbeck Foundations. All genomic data have been deposited to GEO, accession number GSE86906.

## Author Contributions

DCM, SS and NJK prepared the GPCs used in the study, including the cytometry and sorting; SS and LZ performed the transplants; LZ and SS did the histological processing, imaging, and quantitative analyses; JM performed the genomic analyses; MSW and SAG designed the study; and SAG analyzed all data and wrote the manuscript.

